# *DiscoSnp++*: de novo detection of small variants from raw unassembled read set(s)

**DOI:** 10.1101/209965

**Authors:** Pierre Peterlongo, Chloé Riou, Erwan Drezen, Claire Lemaitre

## Abstract

**Motivation:** Next Generation Sequencing (NGS) data provide an unprecedented access to life mechanisms. In particular, these data enable to detect polymorphisms such as SNPs and indels. As these polymorphisms represent a fundamental source of information in agronomy, environment or medicine, their detection in NGS data is now a routine task. The main methods for their prediction usually need a reference genome. However, non-model organisms and highly divergent genomes such as in cancer studies are extensively investigated.

**Results:** We propose *DiscoSnp++,* in which we revisit the *DiscoSnp* algorithm. *DiscoSnp++* is designed for detecting and ranking all kinds of SNPs and small indels from raw read set(s). It outputs files in fasta and VCF formats. In particular, predicted variants can be automatically localized afterwards on a reference genome if available. Its usage is extremely simple and its low resource requirements make it usable on common desktop computers. Results show that *DiscoSnp++* performs better than state-of-the-art methods in terms of computational resources and in terms of results quality. An important novelty is the de novo detection of indels, for which we obtained 99% precision when calling indels on simulated human datasets and 90% recall on high confident indels from the Platinum dataset.

**License:** GNU Affero general public license

**Availability:** <u>https://github.com/GATB/DiscoSnp</u>

## 1 Introduction

Next Generation Sequencing (NGS) data provide an unprecedented access to life mechanisms. In particular, these data enable to assess genetic differences between chromosomes, individuals or species. Such polymorphisms represent a fundamental source of information in many aspects of biology with numerous applications in agronomy, environment or medicine.

With the democratization of sequencing provided by the NGS technologies, determining genetic differences such as SNPs or indels has now become a routine task. There exist numerous software designed for predicting such polymorphisms. Mostly, these methods are based on the use of a reference genome by mapping sequenced reads as this is the case for GATK [6] or SamTools [14] or by mapping partial assemblies as for DISCOVAR [22] or FERMI [12] to cite a few. Basically, they first map the sequences to the reference genome and in a second phase they scan the reference genome to analyse for each locus the differences between the reference sequence and the mapped sequences.

These methods are well accepted and extensively used. However, they present two severe drawbacks. First they are highly sensitive to the mapping quality. Highly repeated regions of the reference genome are difficult to map with a sufficient degree of confidence. Polymorphisms detected from these repeated regions may be spurious as the quantification of mapped reads is erroneous and as the differences between occurrences of the repeats can be wrongly interpreted as polymorphism. Secondly, they suffer from the fact that they need a high quality reference genome. This evident and strong condition limits the application to well studied species. Emerging nanopore sequencing technologies will clearly modify the landscape. However, their current and near future error rate do not break down the barrier of high quality finished assembly and, due to their intrinsic read lengths, they will not provide sufficient coverage for extensible and precise punctual variant calling.

In this context, there is still an important and constant need for *reference-free* methods detecting SNPs and indels, directly from sequenced reads, without requiring any well assembled reference sequence. An alternative method consists in first assemble the reads before to map them back to the so obtained reference, as this is the case in the work by [23]. However, such methods cumulate both the assembly and the mapping difficulties. In this manuscript, we refer to such methods as the *hybrid* strategy.

A few methods [19, 8, 11, 18, 15, 21?] were proposed for *de-novo* detection of polymorphism. All these methods are based on the use of the *de Bruijn graph,* i.e. a directed graph where the set of vertices corresponds to the set of words of length *k*(*k*-mers) contained in the reads, and there is an edge between two *k*-mers if they perfectly overlap on *k*– 1 nucleotides. In this data structure, polymorphisms generate recognizable patterns called *“bubbles”.* These tools detect and analyze such bubbles in order to decipher their origin (sequencing errors, polymorphism due to inexact repeats, real SNP or indel).

In this paper, we present *DiscoSnp++*, an approach that outperforms other *reference-free*methods in terms of skills, of computational needs and of results quality. Its main features are **1**/ the range of predicted variant types: SNPs, isolated or not (an isolated variant is far away from other source of polymorphism), and small indels, **2**/ its extremely low memory usage (several billion reads may be analyzed with no more than 6 GB RAM memory), **3**/ its high execution speed, **4**/ its good precision and recall, **5**/ its faithful score assigned to each predicted variant, **6**/ the output files are in fasta and VCF format, easily usable for downstream analyses and **7**/ the fact that it can be applied to any number of read sets, including only one.

*DiscoSnp++*is a totally new version of *DiscoSnp*[21]. The tool was re-implemented from scratch using the GATB library [7], enabling a much faster running time and a lower memory footprint. The algorithmic models were revisited in order to better filter out sequencing errors and in order to detect new kinds of variants. *DiscoSnp++*outputs the predicted variants in the commonly-used VCF format, optionally after mapping its predictions to a reference genome. This last point may appear counter-intuitive for a reference-free approach, but it is particularly useful when one disposes of a reference genome that can not be used for calling variant by mapping but that can be used for positioning de-novo predicted variants. This can be the case when the reference is either too badly assembled or too far from the sequenced species, which are two very usual situations. Moreover, even in the case of a good reference genome, variant prediction and genotyping with the reference-free approach is not biased in any way by the reference alleles.

## 2 Material and methods

### 2.1 Variant models

#### 2.1.1 Fundamental models

The *DiscoSnp++*tool is based on the analysis of the *de Bruijn Graph* (dBG) [20]. Basically, in a dBG, a *bubble* denotes a path in the graph which diverges into two distinct paths before to meet back. More precisely, we define a branching node as a node that has more that one predecessor and/or more than one successor. A bubble is defined by one first branching node called *start*that has, two distinct right children nodes *n*_*h*_ (higher) and *n*_*l*_ (lower). From *n*_*h*_ and *n*_*l*_, two paths (respectively called *higher* and *lower*) exist and merge in a right branching node. Depending on the kind of variant generating the bubble, the two paths may be of different lengths. If node *start*has more than two right children nodes, all pairs of them are treated as *n*_*h*_ and *n*_*l*_.

In this framework, a variant is called *isolated* if there exists no other polymorphism *k* nucleotides before and *k* nucleotides after the variant position.

In the dBG created from one or several read sets, any isolated SNP generates a simple bubble as shown Figure 1.a. The two paths of such bubbles are composed of exactly *k* nodes each.

**Fig. 1.**
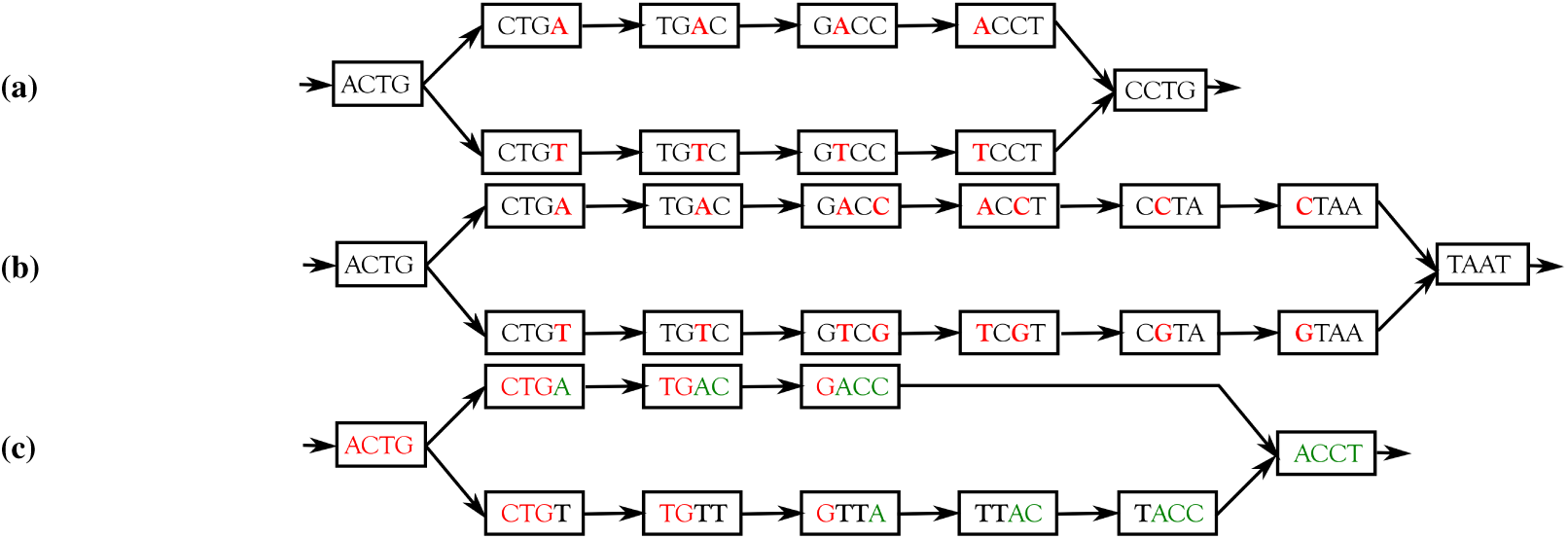
Examples of dBG (k = 4) bubbles due to SNP, close SNPs and indel. (a) Bubble generated by an isolated SNP. Prediction would be ACTGACCTG and ACTGTCCTG. With respect to proposed notations the node ‘ACTG’ is called *start,* the node CTGA is called *n*_*h*_ and the node CTGT is called> *n*_*l*_ (b) Bubble generated by two close SNPs. Prediction would be ACTGACCTAAT and ACTGTCGTAAT. (c) Bubble generated by an insertion. Prediction would be ACTGACCT and ACTGTTACCT.

In the dBG, close SNPs generate a bubble in which the two distinct paths contain the same number of nodes, larger than k. Figure 1.b shows a toy example of such bubble.

As presented in Figure 1.c, indels also generate bubbles in the dBG. The two paths of such bubbles are of different lengths. In theory, indels of any size can be detected. In practice, depending on the data quality and genome complexity, indels longer than a few dozen nucleotides are hardly predicted due to branching paths in the bubble motifs. Note that, however, unlike in mapping approaches, the detected indels are not limited by the read sizes.

The smallest path of an indel bubble is of size at most *k* – 1 nodes. As shown in Figure 2, depending on the indel context, this path may be smaller than *k* – 1. This is the case when the indel and what follows (resp. what precedes) have a common prefix (resp. suffix) of size > 0. Precisely, if the sizes of these common prefix and suffix are respectively *p*_1_ and *p*_2_, the shortest path is composed of max(0, *k* – 1 – *p*_*1*_ – p_2_) nodes and the longest is composed of *k* – 1 + *d* – ***p***_1_ – ***p***_*2*_ with *d* the indel size.

**Fig. 2.**
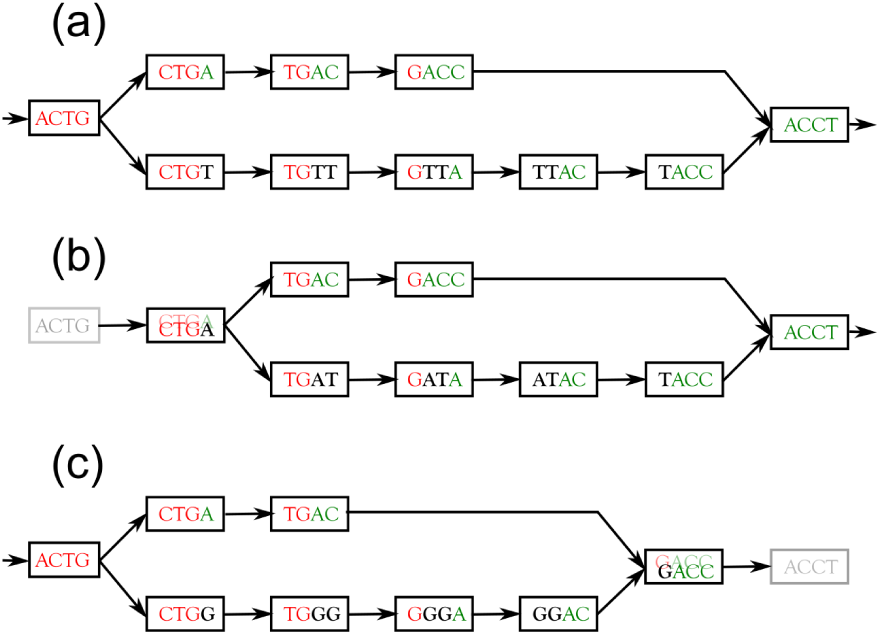
Examples of dBG (*k* = 4) bubbles due to indels with distinct smaller path lengths. Bubble (a) is generated by the presence of sequences ACTGACCT and ACTGTTACCT. The indel is TT. Bubble (b) is generated by sequences ACTGACCT and ACTGATACCT. The indel is AT. In this case, the longest prefix between the indel AT and the following sequence (ACCT) is ‘A’ of size > 0. In this situation, the smallest path is composed by less than *k*-1 nodes. Conversely, in the bubble (c), generated by sequences ACTGACCT and ACTGGGACCT, the indel is GG whose longer suffix with what precede (ACTG) is ‘G’ of size > 0. In this situation, the shortest path is also composed of less than *k*-1 nodes.

Note that in the original *DiscoSnp* approach [21], only isolated SNPs (Figure 1.a) were detectable.

#### 2.1.2 Branching paths

Highly repeated or complex polyploid regions, and sequencing errors present in the dBG yield *“branchings”* in the bubble paths. A bubble is called branching when at least one of its two paths contain a branching node, excepted the rightmost opening branching node and the leftmost closing branching node. All bubbles presented Figures 1 and 2 are non branching.

We differentiate two distinct branching bubble types. The *“non symmetrically”* and the *“symmetrically”* branching bubbles. Intuitively, symmetrically branching bubbles are bubbles in which from nodes *n*_*h*_ and *n*_*l*_, more than a couple of paths can be walked simultaneously. Conversely, in non symmetrically branching bubbles, from those nodes *n*_*h*_ and *n*_*l*_ there exists exactly one unique pair of paths (higher and lower) that merges in a right branching node, whenever those two paths contain branching nodes. Figure 3 presents examples of symmetrically and non symmetrically branching bubbles. Depending on parameters, *DiscoSnp++* detects only non branching bubbles, or non branching and non symmetrically branching bubbles or all king of bubbles without restrictions.

**Fig. 3.**
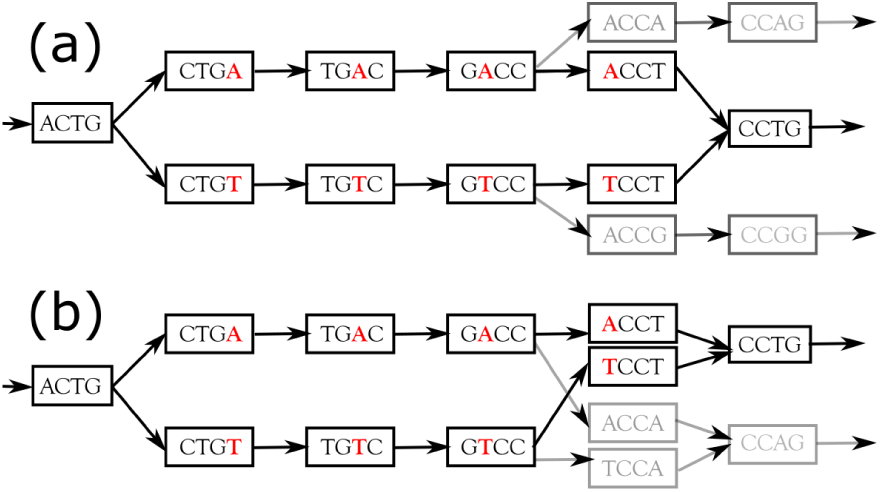
Example of branching and symmetrically branching bubbles. Bubble (a) is branching but non symmetrically branching. From nodes *n*_*h*_ (CTGA) and *n*_*l*_ (CTGT), there exist exactly one path (CCTG) that make the two paths converge, even if some branching nodes are traversed in those paths. Bubble (b) is symmetrically branching. Indeed, upper path and lower path are non unique as nodes GACC and node GTCC can both be extended either with a ‘T’ or with a ‘A’.

### 2.2 Algorithms

This section presents the *DiscoSnp++* main algorithms for detecting bubbles associated to SNPs and indels, once the dBG has been built. The core of the algorithm is to expand conjointly two paths. This is what is done by Algorithm 2. It is called by Algorithm 1, which considers all right branching *k*-mers (with two or more successors) as potential starting *k*-mers of a bubble.

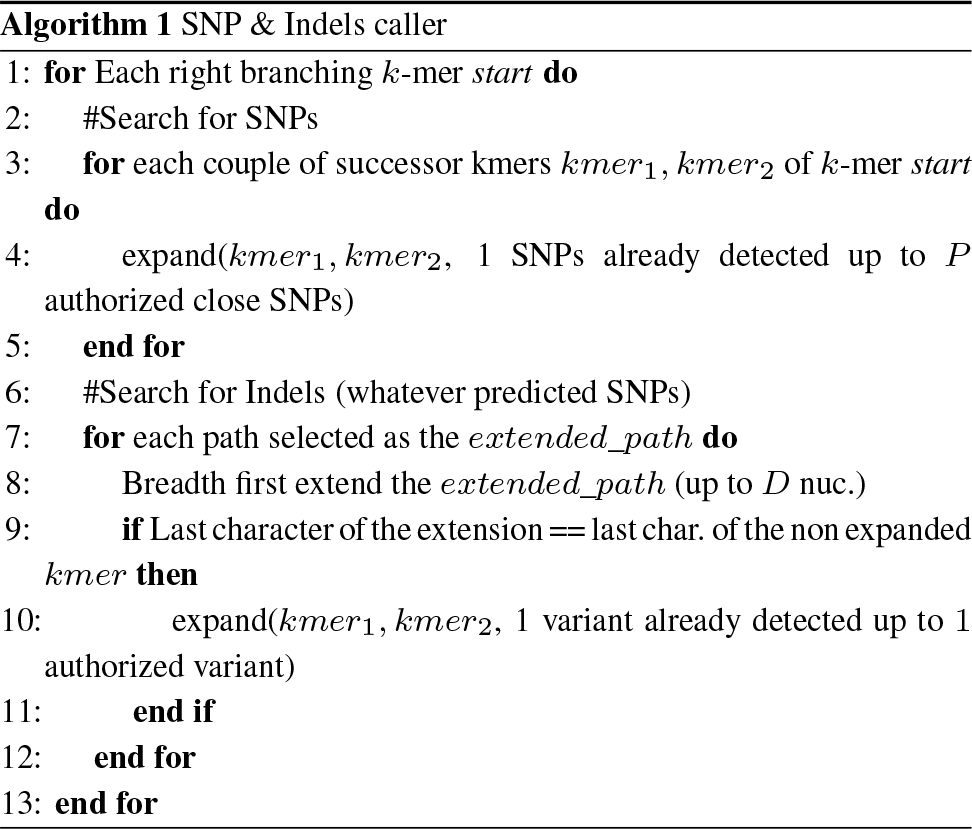

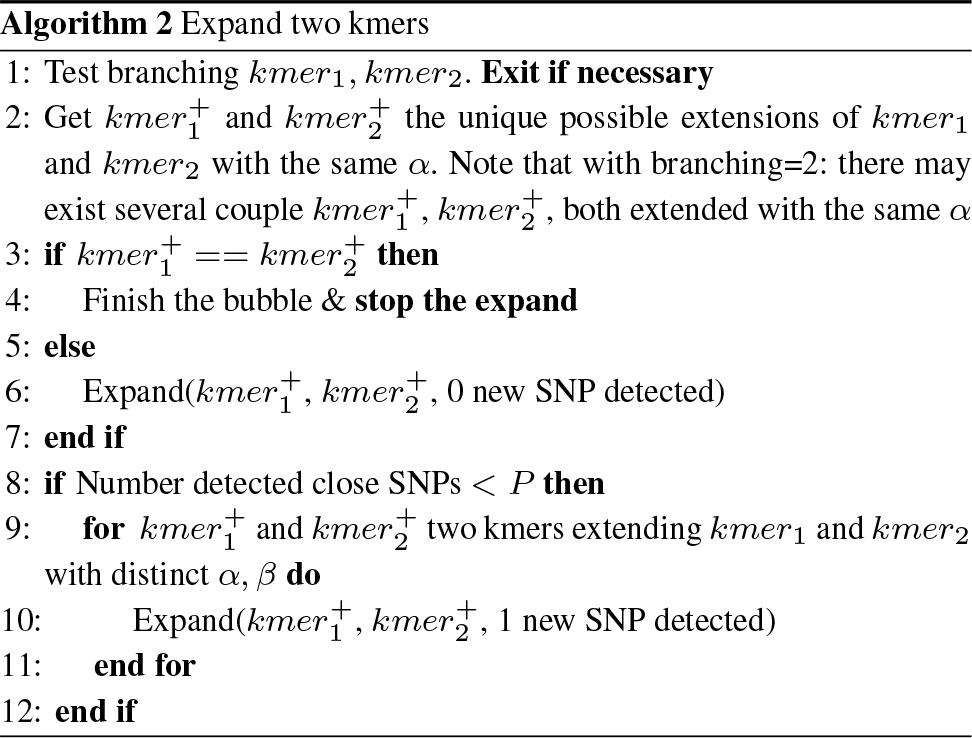

*Details of algorithm 1:* This algorithm traverses all the right branching *k*-mers (for loop in line 1). Each of them is first considered as potentially starting a SNP bubble (lines 3 to 5) and then as potentially starting an indel bubble (lines 6 to 12). In this second case, the algorithm has to propose all possible insertion sequences. Indeed the insertion may be either in the upper or in the lower path (explaining the for loop in line 7), and may be of any size up to the maximal indel size authorized by the user (explaining lines 8 and 9). For instance, in the bubble presented in Figure 2.a, the *extended_path* is the lower one and the breadth first extension was ‘TTA’ whose last character (’A’) is the same than the last one of the *k*-mer *n*_*h*_. In this example, the function *Expand* is called from k-mers ‘CTGA’ and ‘GTTA’.

*Details of algorithm 2:* This recursive algorithm expands conjointly two paths, checking several constraints. First (line 1) it checks if the two *k*-mers to expand respect the constrains on the branching limitations provided by the user. Indeed the user may limit the search to non branching bubbles, to non branching or non symmetrically branching ones, or to any kind of bubbles. In the case of non branching or non symmetrically branching bubbles there must exist only one unique character (A,C,G or T) that can expand the two *k*-mers. For simplifying the reading, the case of symmetrically branching nodes is not presented here. In practice, this latter case consists in considering several couples 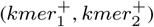 extending the input *k*-mers instead of only one. If the two extending *k*-mers are identical, the bubble finishes (lines 3 and 4), else the recursion continues. Whatever the result of expansion, the algorithm checks the existence of potential close SNPs (lines 8 to 11) when input k-mers *kmer*_*1*_ and *kmer*_*2*_ can be extended with different characters α and β.

#### 2.2.1 Reverse complement and canonical bubbles

The strand of raw reads is unknown. Thus each indexed *k*-mer can be read either in the forward strand or in the reverse complement strand. Consequently each bubble is found once from left to right and once from right to left. For instance topological bubble from Figure 1, generates the sequence couple “ACTGACCTG” and “ACTGTCCTG” and the other sequence couple “CAGGTCAGT” and “CAGGACAGT” when read from right to left. For avoiding the redundancies, only one of the two versions, called *“canonical”,* is output. The canonical version of a variant is the version which contains the lexicographically smaller sequence. In the previous example, the couple “ACTGACCTG” and “ACTGTCCTG” is the canonical representation of the variant. Note that, the bubble has to be fully detected to know if it is canonical or not as the canonicity depends on the entire sequences. This doubles the computation time and this could be avoided at the expense of a higher memory footprint by marking *k*-mers already used in a previously detected bubble.

### 2.3 Implementation

#### 2.3.1 Pipeline

**Fig. 4.**
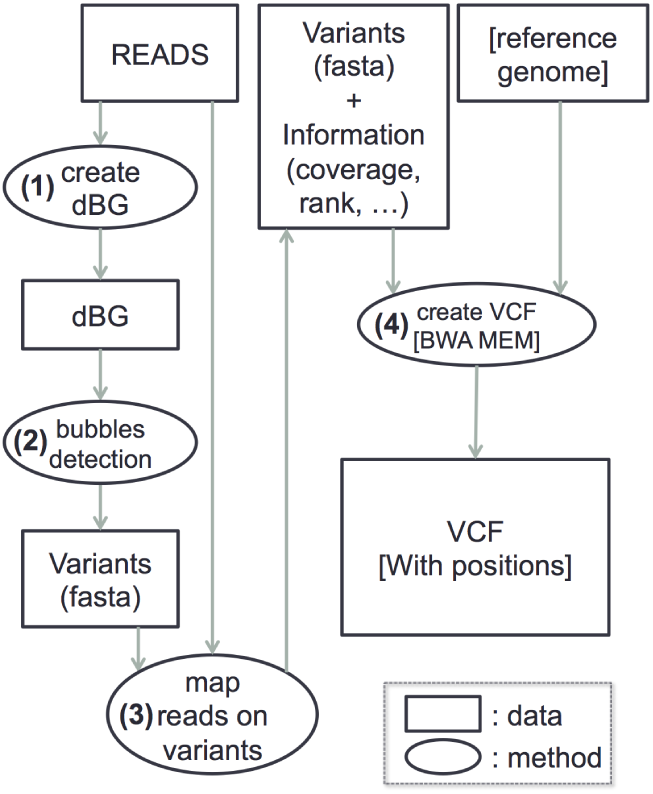
*DiscoSnp++* pipeline. If a reference genome is provided then VCF_Creator uses BWA-mem [13] and outputs a VCF file that contains the localization of mapped predictions.

*DiscoSnp++* is composed of several independent tools. The pipeline is presented in Figure 4.

(**1**) The dBG of the input dataset(s) is built. This step relies on the GATB library [7]. This step includes notably the removal of erroneous *k*-mers (see Section 2.3.2). (**2**) a module detects bubbles generated by the presence of SNPs (close or not) and indels; This module implements Algorithms 1 and 2. It outputs a fasta file containing the variant sequences together with the contig (or unitig or nothing, depending on the user choice) they belong to (see Section 2.3.3). (**3**) For keeping the RAM usage very low, no other information than *k*-mer presence is stored in the dBG. Thus the variant sequences do not contain any information about abundances in read dataset(s). A module maps back the reads from all input read sets on the variant sequences, mainly in order to determine the read coverage per allele and per read set of each variant. This module also computes a rank per variant, indicating whether the variant shows discriminant allele frequencies between compared datasets. When two datasets are compared, this rank is the Phi coefficient of the contingency table of read counts, computed as follows: 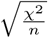, with *n* being the sum of read counts. When more than two datasets are compared, the rank is the maximal Phi coefficient obtained among all possible pairs of considered datasets. When two or more read sets are used, this score offers a way to filter out putative false positive variants due to inexact repeats as they are likely to have similar allele frequencies in all datasets. Optionally, as presented Section 2.3.4, a genotype per read set is computed for each predicted variant. (4) The last module generates a VCF of the predicted variants. If no reference genome is provided, this step is simply a change of format from fasta to VCF. If a reference genome is provided, this module maps predicted variants on this reference using BWA-mem [13]. This enables to output for each variant its localization in the genome. If a variant has more than one best mapping position, then one of those positions is randomly output, and a VCF field indicates a multiple mapping. A variant with multiple mapping positions is likely to be due to inexact repeats and numerous false positive variants can be easily filtered out using this information. The detailed VCF content is described in the additional file.

#### 2.3.2 Filtering out sequencing errors

Raw NGS reads contain sequencing errors, estimated between 0.1% and 1%. Each sequencing error generates up to *k* erroneous *k*-mers. A crucial step consists in removing those *k*-mers from the dBG. This is done using a classical *k*-mer counting approach using the GATB library, implementing the KMC2 algorithm [5]. Then, *k*-mers whose occurrence number is below a solidity threshold *T*_*sol*_ are removed. In *DiscoSnp++,* a solidity threshold is defined independently for each read set (as read sets may have been obtained with different sequencing efforts). A *k*-mer whose occurrence number is higher than the solidity threshold for any of the read sets is considered as solid and is conserved in the dBG, otherwise, it is discarded. For each read set, the optimal T*soi* value can be inferred automatically from the analysis of the *k*-mer counts profile, with a method similar to the one used in kmergenie [3].

More precisely, the T*soi* parameter is inferred automatically from the histogram of *k*-mer abundances (telling the number of kmers of each abundance). The following heuristic is used. The histogram is first smoothed, then we search for the position of the first increase, and the maximum value attained after that. We then look for the index of the minimum value between this first raise and this maximum. This index is the automatically inferred threshold, and roughly corresponds to the valley between erroneous kmers and solid kmers. We also ensure that no more than 25 % of the distinct kmers are below the threshold, and that the threshold is ≥ 3.

*DiscoSnp++* implementation enables for each read set to manually fix a solidity threshold or to use the automatic detection.

#### 2.3.3 Assembly of context contigs

Depending on the user choice, the contig or unitig located on the left and the one right of the predicted bubble are computed. A unitig is a simple non branching path of the dBG and a contig, as defined in this framework, is a walk in the dBG that may crush some small bubbles. This step uses the Minia [4] assembler implementation.

#### 2.3.4 Predicting genotypes

For each prediction and each dataset, a genotype is provided assuming each dataset corresponds to a diploid individual. Genotypes are inferred independently for each individual, based on the read coverage of each allele of the bubble. To do so, the likelihoods of the three possible genotypes (homozygous 0/0 or 1/1 or heterozygous, 0/1) are computed based on a simple binomial model as described in the Nielsen 2011 review [17]:

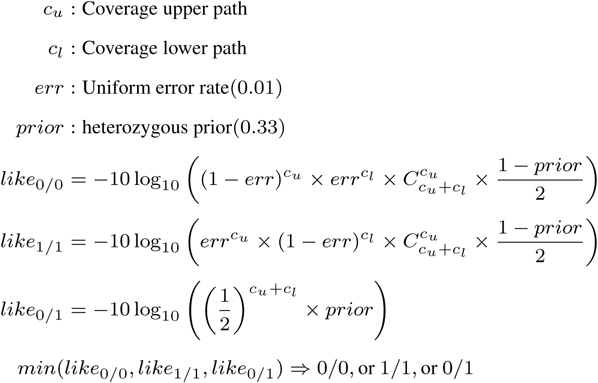

These computations rely namely on the *error probability* parameter, that is the probability that a read maps to a given allele erroneously, assuming it is fixed and indepentent between all observations. It was fixed to 0.01, referring to classical sequencing error rates.

Finally, the genotype showing the largest likelihood is chosen and all three likelihoods are also output (-log10 transformed) as additional information. Notably, only the probability of observing the data (ie. the read counts for each allele) given the genotype is computed, no prior is used so as to compute the posterior probabilities of each genotype given the data. Users may deactivate genotyping, for instance when the input datasets are not coming from diploid individuals. For a given read set, if a variant has a read coverage lower than the solidity threshold (see Section 2.3.2) for both alleles, its genotype is annotated as missing and denoted by “.|.”.

#### 2.3.5 Practical read sets representation

We differentiate *“read files”* from *“read sets”.* A read set may be composed of two or more read files, and the *DiscoSnp++* input are read sets. This feature is particularly important when a given sequenced sample is spread over two files (for instance with paired-end or mate pair sequencing) or more (sequencing lanes, replicates, pooling, or any other reason).

Our implementation allows the use of a reference genome as an input set. This enables to predict homozygous variants having the same allele in all read sets distinct from the allele in the reference genome.

#### 2.3.6 Default parameters

By default *DiscoSnp++* uses the automatic solidity threshold detection, *k*-mers of size 31 and predicts variants from non branching bubbles and non symmetrically branching bubbles. By default, at most 10 SNPs per bubble and indels of size at most 100 are searched.

### 2.4 Materials

#### 2.4.1 Simulation of 30 haploid bacterial individuals

We propose an experiment simulating more than two haploid bacterial individuals. For doing this, we created 30 copies (that we call individuals) of the *E. coli K-12 MG1655* strain. We then simulated SNPs with a uniform distribution such that ≈ 4,200 SNPs (≈0.1% of the genome length) are common to any pair of individuals, half this number is common to any trio of individuals, a third to any quadruplet, and so on. With this strategy, when considering all the 30 individuals together, 69,600 SNP sites were generated, covering ≈1.5% of the genome. We also simulated indels following exactly the same process, with a ratio of one indel for ten SNPs.

We simulated a 40x sequencing of each of the 30 individuals, with 100-bp reads and 0.1% error rate. Thus, 1,855,870 reads were generated per read set.

*DiscoSnp++* was run on the two first read sets, then on the three first read sets, and so on up to the whole 30 read sets. Each computation was performed using the default parameters, looking for indels of size at most 10 and up to four close SNPs per bubble.

All used commands for all tested methods can be found in the Additional File.

#### 2.4.2 Simulation of two diploid human individuals (limited to chromosome 1)

We propose an experiment simulating two human diploid read sets. This experiment applied on human chromosome 1, (GRCh37/hg19 reference assembly version), ≈ 249 million base pairs. SNPs and indels were simulated following the 1000 genome project [2] predictions. Precisely, from the phase 1 vcf file [1], we extracted SNPs and indels of individuals HG00096 and HG00100. Variants having the same homozygous genotype in both individuals were discarded. For each of the two individuals, two versions of the chromosome were created. On each of these two versions, SNPs and indels were placed following the VCF file, i.e., 0|0: no modification of the chromosomes, 0|1: modification of one of the two chromosomes (randomly chosen), 1|1: modification of the two chromosomes taking into account the phase information.

For each individual, we simulated a 40x illumina sequencing with 0.1% error rate (20x per chromosome). Thus, 99,600,000 reads were generated per read set. *DiscoSnp++* was run on the so obtained two read sets, using default parameters, looking for indels of size at most 10 bp and up to four close SNPs per bubble.

Again, all used commands for all tested methods can be found in the Additional File.

#### 2.4.3 Real data, saccharomyces cerevisiae experiment

We used a set of biologically validated variants predicted from an artificial evolution study on *Saccharomyces cerevisiae* [9]. In this study, three glucose-limited, chemostat-evolved populations of the haploid yeast strain S288c, named E1, E2 and E3, were sequenced every ≈ 70 generations, giving eight samples per population. Using a reference-based mapping approach, 110 mutations were discovered, among which only 33 have a minor allele frequency (MAF) > 10% and 32 were confirmed by Sanger sequencing.

*DiscoSnp++* was run independently on populations E1, E2 and E3 (reads downloaded from the NCBI Sequence Read Archive, accession number SRA054922). For each population, *DiscoSnp++* was applied on the eight read sets corresponding to the eight time points, with the default parameters and T_sol_ = 11, D = 60 (searching for indels of length at most 60 bp) and *P* = 4 (authorizing up to 4 close SNPs in a unique bubble).

#### 2.4.4 Real data, human platinum

We used reads from the Platinum study [?]. More precisely, we downloaded the reads of individuals NA12877 and NA12878, sequenced with 200x coverage depth on a HitSeq 2000 system. These two datasets each benefit from a VCF file^1^ containing variants called by a set of nine independent methods. We call those files NA12877.vcf and NA12878.vcf. Thus each variant may be found by any number of methods between one and nine.

We ran *DiscoSnp++* on the two read sets, using the human genome (ref. hg19) for mapping predictions and also as an input set for predicting variants distinct from the reference but identical in the two sets.

We validated the predictions by comparing the VCF generated by *DiscoSnp++* with the two VCF files NA12877.vcf and NA12878.vcf limited to variants found by at least half of the methods employed in the platinum study.

#### 2.4.5 Results validation

In all the performed tests, we analyzed the VCF file containing predicted variants of *DiscoSnp++* localized afterwards on the reference genome. We compared it to a reference VCF, containing either the simulated variants (ground truth) or variants validated by other methods (for real data). Within this approach the number of True Positives (TPs): correctly predicted variant; of False Positives (FPs): wrongly predicted variant; of False Negatives (FNs): real variant, non predicted, can be easily computed.

Precision of a prediction set is then defined by

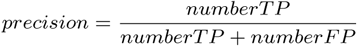

and the recall is defined by

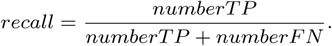

Note that in the case of real datasets, the *precision* is meaningless as the reference sets are not exhaustive. The main manuscript does not provide *precision* for real datasets.

## 3 Results

We propose results both on synthetic and on real datasets. Synthetic datasets offer a way to exactly compute the precision and the recall of *DiscoSnp++* and of state-of-the-art methods.

### 3.1 Results on synthetic datasets

We first propose a bench of results based on synthetic datasets. These datasets are derived from real genomes, either from *Escherichia Coli* or from the human chromosome 1.

We tested *DiscoSnp++, cortex* [8] and an hybrid method composed of SOAPdenovo2 [16] for generating the assembly, Bowtie2 [10] for mapping the reads on the assembly, and GATK [6] for calling variants. Presented results were obtained using *UnifiedGenotyper* option of GATK. As presented in the Additional File, when following the GATK guidelines (including read realignment and using the *HaplotypeCaller* option), the result quality is similar but at the expense of a much longer execution time (almost 3 times longer, from approximately 19h to 54h for the human chromosome 1 experiment).

#### 3.1.1 TWO and more bacterial read sets

We performed an experiment on a variable number of read sets. Each read set corresponds to a simulation of the sequencing of an *Escherichia Coli* individual.

Precision and recall results, presented in Figure 5, allow to demonstrate that *DiscoSnp++* is the only tool able to provide high quality results (recall and precision) both for SNPs and indels and for large number of haploid individuals.

**Fig. 5.**
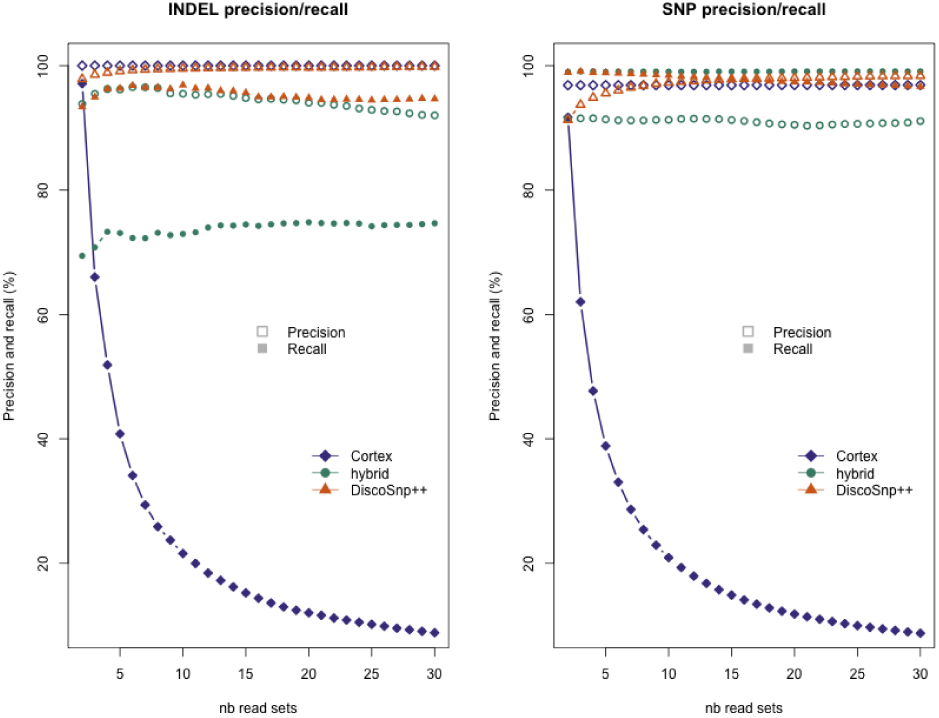
*DiscoSnp++*, cortex and hybrid strategy (SOAPdenovo2+GATK) results, depending on the number of input haploid individuals. SOAPdenovo2 and GATK were launched with default parameters. For *DiscoSnp++* and cortex, *k*-mers having three or fewer occurrences in all datasets were removed. Precision and recall: filled symbols represent the precision and empty symbols represent the recall. Left: results on indels predictions. Right: results on SNPs predictions.

In more details we draw the following conclusions. On these data, for calling both SNPs and indels, the *cortex* precision is perfect or nearly perfect, whereas its recall strongly decreases when the number of read sets increases, reaching less than 9% for 30 individuals.

For SNP calling, the hybrid and *DiscoSnp++* approaches provide excellent recall (respectively at least 98.98% and 96.65%). We may however note that recall slightly decreases for *DiscoSnp++* when increasing the number of read sets whereas this is not the case for the hybrid approach. The precision is stable for the hybrid approach whatever the number of read sets (≈ 91%), while it increases from 91% to 98% with the number of read sets for the *DiscoSnp++* approach.

Concerning indels, the hybrid strategy shows bad performances, that may be explained by the difficulty of mapping reads with indels. Conversely, *DiscoSnp++* presents high quality results in terms of both precision and recall.

In short, in terms of results quality, *DiscoSnp++* outperforms the compared methods, with a major advantage in indel calling. Importantly, as shown in Figure 6, *DiscoSnp++* runs much faster that other methods and uses much less RAM memory. Moreover, one may also insist on the fact that *DiscoSnp++* is extremely simple to use. Table 1, showing the number of operations to perform for each method, witnesses unequivocally this simplicity.

**Table 1.** Complexity in term of required number of command lines (including file formatting when necessary) while calling variants from 2 and 30 haploid genomes.

|  | Number of<br>command lines<br>for two genomes | Number of<br>command lines<br>for 30 genomes |
| --- | --- | --- |
| Hybrid | 19 | 187 |
| *cortex* | 8 (+2 compilations) | 35 (+30 compilations) |
| *DiscoSnp++* | 2 | 2 |

**Fig. 6.**
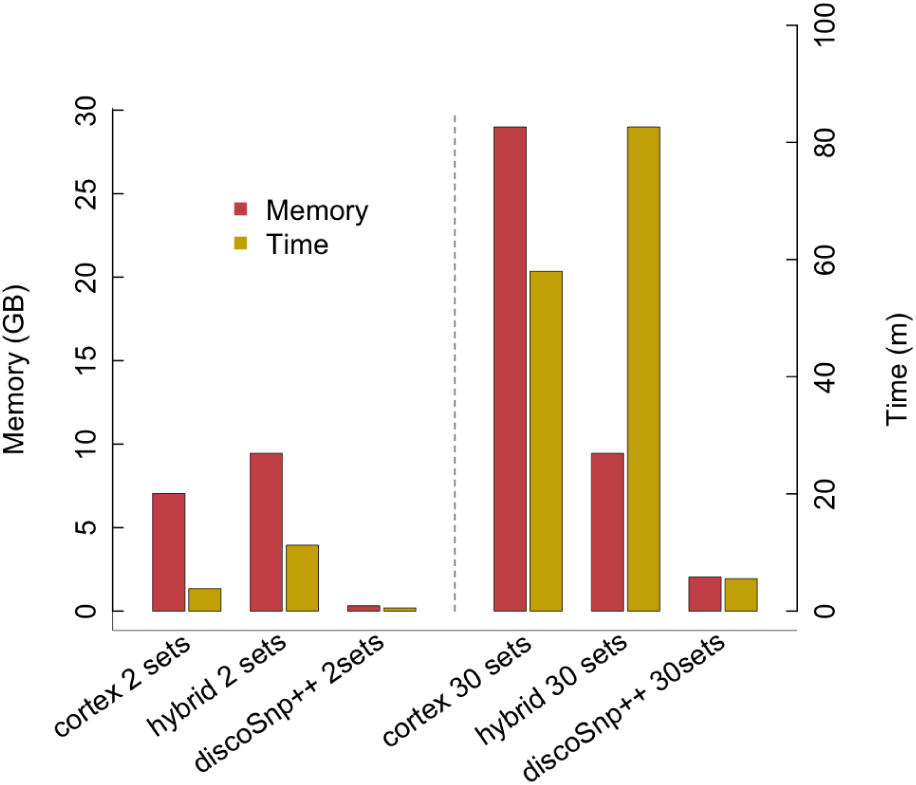
Comparative memory and time performances, depending on the number of input haploid individuals.

### 3.1.2 TWO human datasets

To evaluate *DiscoSnp++* on a diploid and more complex genome, we propose an experiment based on the human chromosome 1, assembly GRCh37. Using variants from the 1000 genomes project, we simulated two individuals, generating 25,928 indels in the range 1 to 10 bp (average 2.1 bp) and 288,069 SNPs.

Results considering all predictions are presented Table 2. Precision and recall of SNP calling are quite similar between compared tools, with *DiscoSnp++* achieving the best recall and overall the best compromise between both values. Surprisingly, tools show more contrasted results on indels. In particular, *DiscoSnp++* finds almost twice as much true indels than the hybrid approach (72 vs 41 % recall). This comes with a slightly lower precision, but filtering *DiscoSnp++* predictions on the basis of the rank score enables to reach a much better precision without losing too much in recall. For instance, indels with a rank higher than 0.2 are almost all true ones (99.25% precision) with a recall still much higher than the one of the hybrid approach (58 %).

**Table 2.** Human chromosome 1 results. “Rec”. stands for “Recall” and “Prec.” stands for “Precision”

|  | SNP |  | indels |  |
| --- | --- | --- | --- | --- |
|  | Prec. (%) | Rec. (%) | Prec. (%) | Rec. (%) |
| Hybrid | 71.60 | 78.59 | 96.15 | 40.97 |
| *cortex* | 73.19 | 67.34 | 86.65 | 63.25 |
| *DiscoSnp++* | 72.11 | **79.31** | 77.49 | **71.74** |
| *DiscoSnp++* (rank > 0.2) | **98.73** | 65.17 | **99.25** | 57.59 |

Results presented Figure 7 provide additional pieces of information for the hybrid and the *DiscoSnp++* approaches. They show precision/recall values with respect to the ranking of the predictions *(cortex* results are not ranked in this framework). Results show that the hybrid approach predictions are badly ranked: it appears that predictions showing the best scores are mainly false positives. Additionally, results show that the *DiscoSnp++* ranking is extremely efficient for distinguishing false positives from true positives. Most of the predictions ranked with a score > 0.2 are true positives (98.73% of the SNPs and 99.25% of the indels).

From these experiments on a complex eukaryotic species, one may get the feeling that *DiscoSnp++* overall results are similar to the ones from other methods. However, it is the only tool with a reliable ranking of the results, enabling to select more than 50% of the predictions with a nearly perfect precision when filtering results based on its computed ranking.

The low recall is mainly explained by too factors. First, complex and repeated regions generate symmetrically branching bubbles that were not predicted in this experiments. When we include symmetrically branching bubbles, the recall raises to 95.36% (with a precision of 11.15%). Importantly, this reflects the fact that most missed variants fall in complex repeated genomic regions that generate symmetrically branching bubbles. Second, the filtering of erroneous *k*-mers is not perfect, and it can remove a few non erroneous *k*-mers. Finally, including symmetrically branching bubbles and keeping all *k*-mers raises the recall to 99.06% (with a laughable precision of 0.09%). These results show that the *k*-mer filtration is a critical step for avoiding false calls.

**Fig. 7.**
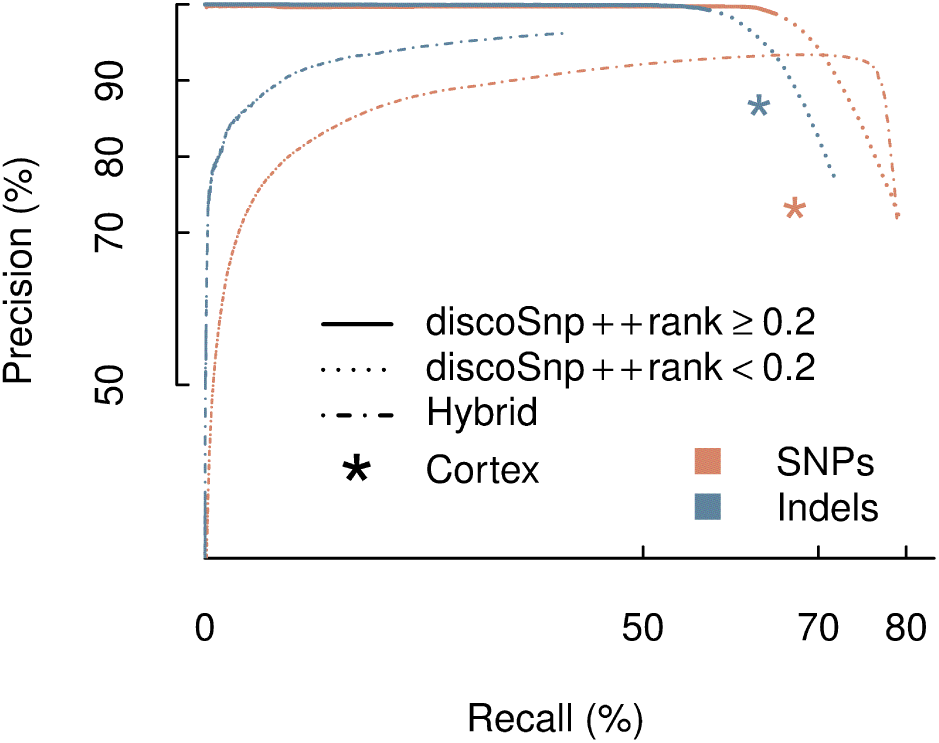
Comparative results of *DiscoSnp++*, cortex, and the hybrid SOAPdenovo2 + Bowtie2 + GATK approaches on the two diploid human chromosome 1 dataset. Precision versus recall curves are obtained by ranking the predicted SNPs and indels. Each data point is obtained at a given rank threshold, where precision and recall values are computed for all SNPs with better ranks than this threshold. The dashed tail of the two *DiscoSnp++* curves denotes the predictions ranked with a threshold bellow 0:2. In this framework *cortex* does not rank its predictions, its results are thus represented by a single point.

As previously mentioned, *DiscoSnp++* provides for each predicted variant, its genotype estimation for each individual. On this dataset, 247,065 true positive variants were predicted thus generating 494,130 predictions (one per individual). Over those predicted genotypes, 485,673 (98.29%) were correct.

When mapping results on a reference genome, among other pieces of information, the number of mapping positions is given for each variant in the vcf file. Considering only variants having a unique mapping position in the reference genome enables to select those that are less likely to be due to inexact repeats. As shown Table 3, this is verified in this dataset as the precision raises from 72.1 % to 87.6% when considering onlu uniquely mapped SNPs.

**Table 3.** Human chromosome 1 *DiscoSnp++* results (number of TP and FP) with respect to the type of variant (SNP or indel) and to the number of mapping positions on the reference genome. Non mapped variants can not be classified as TP or FP.

|  | SNP |  | indels |  |
| --- | --- | --- | --- | --- |
|  | TP | FP | TP | FP |
| Unique | 233,483 | 33,111 | 10,086 | 1,075 |
| Multiple | 3,920 | 45,221 | 174 | 1,501 |
| None |  | 1,097 |  | 0 |

As shown in Figure 8, *DiscoSnp++* runs much faster that other methods (3.6x and 17.5x times faster than *cortex* and the hybrid approach respectively) and uses much less RAM memory (36.2x and 22.9x times less memory than *cortex* and the hybrid approach respectively).

**Fig. 8.**
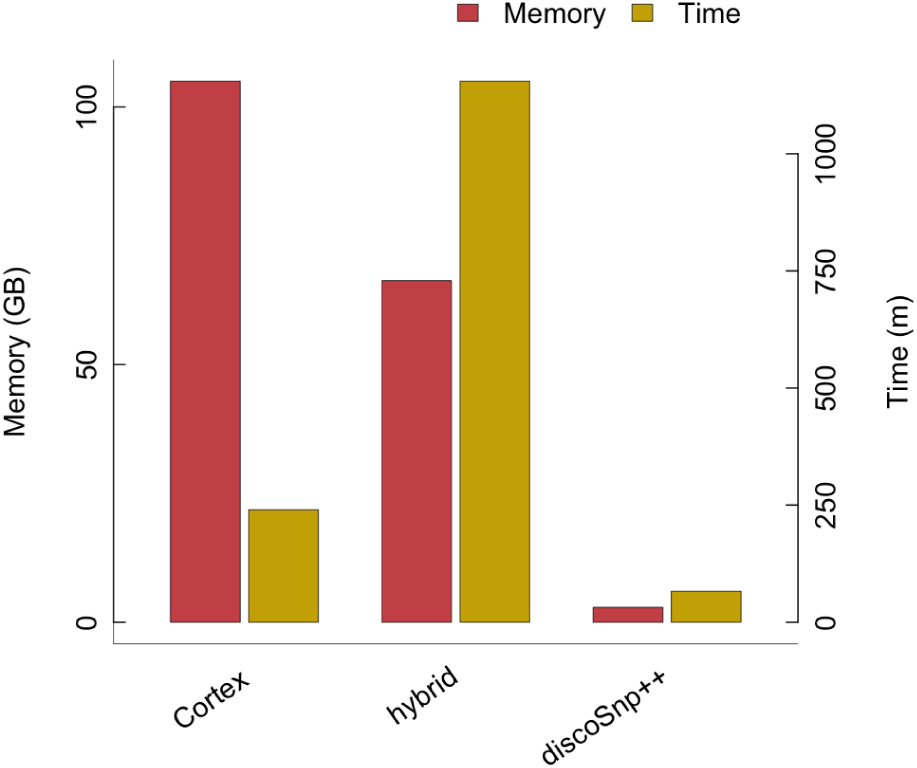
Comparative memory and time performances for comparing the two human datasets

### 3.2 Results on real datasets

#### 3.2.1 Saccharomyces cerevisiae experiment

This experiment aims to demonstrate the recall permitted by *DiscoSnp++* on an experiment where a subset of called variants have been biologically validated.

Among the 32 experimentally validated SNPs, 29 were predicted by *DiscoSnp++,* leading to an estimated recall of 90.7%. Among those 29 SNPs, four SNPs were close.

Detecting all bubbles including symmetrically branching ones leads to the detection of the three remaining unpredicted SNPs. The fact that these SNPs were not detected with the default parameter (detection of non branching and non symmetrically branching bubbles) suggests that these SNPs are located in complex regions of the genome.

Note that in the study from [9], no SNP with a MAF < 10% was validated and no indel was validated, so we could not assess the recall of these types of *DiscoSnp++* predictions on this dataset.

Finally, all indels predicted in the original study except one were also predicted by *DiscoSnp++*, including in particular an indel of size 56 bp.

#### 3.2.2 Platinum experiment

We ran *DiscoSnp++* on two real datasets from two human individuals, NA12877 and NA12878, summing to 400x coverage. The whole process took 22h and required only two command lines.

We compared results obtained by *DiscoSnp++* with those obtained in the Platinum study, found by at least half of the used methods, that is to say at least five methods. This represents a total of 3,174,898 SNPs and 148,194 indels. *DiscoSnp++* predictions retrieved 2,605,955 (82.08%) of those SNPs, and 133,994 (90.42%) of those indels.

When limiting Platinum variants to those validated by at least six methods among nine, 1,714,222 SNPs remain. *DiscoSnp++* found 1,681,626 (98.10%) of them. Interestingly, no Platinum indel are predicted by at least six methods. This highlights the lack of methods for predicting indels with a high accuracy, even among the reference-based methods. In this context we can not conclude about the precision of the 360,798 indels predicted by *DiscoSnp++.* However, it is important to remark that in the context where a reference genome is available, our approach first predicts indels before to map them on the reference. Since for each predicted variant, one of the alleles does not contain the indel, it is easy to map them on the reference. This way, contrary to read mapping approaches, *DiscoSnp++* does not suffer from split mapping issues for predicting indel variants or for finding their genomic locus.

## 4 Conclusion

Small variant prediction remains a fundamental task that is usually a preliminary step, vital to most downstream bioanalyses on sequenced data. Despite the sequencing “democratization” and the impressive current technological improvements, when working on complex species such as eukaryotic ones, there is still only a few reference species for which one disposes of an assembled genome, good enough to serve as support for predicting efficiently and extensively variants of interests.

At the same time, there were still an important lack of user-friendly reference-free variant callers able to scale-up large datasets and to provide results quality similar to or better than reference based approaches. With this work we hope we bridged this gap with *DiscoSnp++,* a new easy to use tool, using low resources and offering high quality results.

The advantages of *DiscoSnp++* over other reference-free methods is even more substantial for indels and our results tend to suggest that for these variants reference-free approaches may play a crucial role even when a reference genome is available.

All variant prediction methods, with or without a reference genome, face limitations mainly in highly repeated genomic regions. Variants embedded in such regions are difficult to predict, preventing from obtaining high recall results. An open problem consists in developing new methods for finding such variants, nowadays inaccessible with a good precision. This could be realized by integrating long Pacbio or Nanopore reads and working on a A-Bruijn graph [20] instead of a de-Bruijn graph. This would resolve repeats smaller than long reads and would allow to extricate variants from their repeated regions.

## Footnotes

1 ftp://ussd-ftp.illumina.com/older_releases/hg19/IlluminaPlatinumGenomes_v7.0/merged_platinum/

## 5 Acknowledgements

We thank the *GenOuest* (genouest.org) cluster team, who allowed us to perform all the tests. This work was supported by the French ANR-12-BS02-0008 *Colib’read* project and by the ANR-12-EMMA-0019-01 GATB project.

